# Facilitating User Interaction with the Tuberculosis Mutation Catalogue using AI Tools

**DOI:** 10.1101/2025.04.25.650567

**Authors:** Miguel Moreno-Molina, Anita Suresh, Rebecca E. Colman, Timothy C. Rodwell

## Abstract

The World Health Organization (WHO) 2023 Mutation Catalogue for *Mycobacterium tuberculosis* is a crucial knowledgebase and tool for clinical interpretation of mutations associated with drug-resistant TB. However, the document’s complexity and size pose challenges for many users. This study evaluated the potential of generative artificial intelligence (AI) models to facilitate natural language user interaction with the catalogue. Four prominent AI models—Google Gemini 2.5 Pro, OpenAI ChatGPT 4.1, Perplexity AI, and DeepSeek R1—were assessed through general test questions, mutation search and retrieval tasks using both full catalogue queries and antibiotic-specific tables, and the application of additional grading rules to score novel mutations. Performance was measured based on accuracy, completeness, clarity, source citation, and the presence of hallucinations. Google Gemini 2.5 Pro consistently demonstrated superior performance in accuracy, completeness, and avoidance of hallucinations across most evaluations, especially in general queries and large dataset searches. DeepSeek R1 excelled in applying grading rules to novel mutations and showed high accuracy in focused datasets, but exhibited some hallucinations. ChatGPT 4.1 was strong in clarity but lacked proper citations, and Perplexity AI showed variable performance with a higher frequency of hallucinations. The findings highlight the potential of AI tools to enhance accessibility and utility of complex knowledgebases like the WHO Mutation Catalogue, while emphasizing the critical need for careful model selection and rigorous benchmarking to ensure accuracy and reliability. The results suggest that Google Gemini 2.5 Pro and DeepSeek R1 are strong candidates for developing a custom clinical AI agent to assist all levels of healthcare users in navigating and interpreting the complex information within the catalogue, ultimately contributing to improved TB control efforts.

## INTRODUCTION

The 2023 Mutation Catalogue for tuberculosis (TB) published by the World Health Organization (WHO), is a critical resource in the fight against drug-resistant tuberculosis (DR-TB) [1]. It serves as the largest global repository of mutations associated with drug resistance in *Mycobacterium tuberculosis*, the bacterium that causes TB. The catalogue offers valuable insights into the genetic basis of drug resistance and clinical interpretation of mutations, which in turn inform treatment regimens, development of new diagnostic tools, and public health strategies. The target audience for this catalogue comprises diagnostic developers, researchers, clinicians, and public health officials involved in TB control and research.

Drug resistance is a major challenge in TB control, as it can lead to treatment failure, increased transmission, prolonged illness and significantly higher mortality [2]. By cataloging *M. tuberculosis* genetic mutations associated with clinically relevant phenotypic resistance to various anti-TB drugs, the WHO catalogue enables public health researchers to track the emergence and spread of drug-resistant strains, diagnostics developers to identify new resistance targets for novel assays, and clinicians to improve the speed and adequacy of their treatment decisions.

Despite its importance, the 2023 WHO mutation catalogue presents a few challenges for many of its intended users. The complexity of the information, coupled with the large size of the dataset, can make it difficult for audiences to navigate and extract relevant information efficiently. Searching for and retrieving specific information can be cumbersome, and systematically applying the catalogue to classify mutations from sequencing data requires specialized bioinformatics expertise. These challenges can hinder the accessibility and utility of the catalogue for a broader audience, including those without extensive expertise in the TB genomics field [3].

To address these challenges and improve user experience, natural language artificial intelligence (AI) tools present a potential solution. Generative AI tools, with their ability to process and analyze large datasets, can potentially facilitate user interaction with the 2023 Mutation Catalogue by providing a more intuitive and user-friendly interface. These tools can potentially simplify the search and retrieval process, assist in data interpretation, and provide tailored information—within the bounds of the training material—to users with varying levels of expertise. While the potential of AI in biomedical research is increasingly recognized[4], systematic evaluations of large language models (LLMs) for natural language interaction with complex, specialized knowledgebases like the WHO Mutation Catalogue remain extremely limited. This study provides a direct comparison of current leading LLMs on specific tasks relevant to catalogue users. The specific objectives of this study were to evaluate the ability of different AI models to accurately interrogate the 2023 WHO Mutation Catalogue, assess their potential to provide correct answers without generating false information (hallucination) or providing information not contained within the source catalogue. We also tested search and retrieval capabilities of each model, and finally, evaluated their understanding and application of the additional grading rules when presented with novel mutations.

## METHODS

### Selection of AI models and input material

The AI models chosen for this study were Google Gemini 2.5 Pro, OpenAI ChatGPT 4.1, Perplexity AI and DeepSeek R1. These models were selected based on a variety of factors, and their widespread availability and accessibility ensured that the research could be conducted without undue constraints. We discarded models such as Microsoft Copilot or Claude 3.5 Sonnet due to their inability to ingest the entire catalogue document as context. The diversity in architectures of the selected models and their training methodologies provided a broad spectrum for analysis, enabling a more comprehensive understanding of the capabilities and limitations of different AI approaches.

Furthermore, these models have all demonstrated significant proficiency in natural language processing (NLP) and information retrieval tasks, which are central to the objectives of this study. Their ability to understand, interpret, and generate human-like text, as well as to search and retrieve relevant information from datasets, makes them well-suited for the demands of this research. Additionally, the selection of these models reflects a deliberate balance between commercial and open-source options. This diversity allows for a more nuanced assessment of the AI landscape and a wider array of future options for implementation. By incorporating a range of AI models with different strengths, weaknesses, and underlying philosophies, this study aimed to provide a holistic and insightful perspective on the potential of AI for this specific application, as well as for the broader field of AI research and development.

For each evaluation (Prompt 1, 2, and 3), the respective AI model was provided with the specific prompt text along with the relevant document file as context. For Prompts 1 and 3, and the full catalogue query testing in Prompt 2, the input file was the complete second edition of the 2023 WHO Mutation Catalogue PDF (specific file available at https://www.who.int/publications/i/item/9789240082410, total of 140 pages). For the antibiotic-specific table query testing in Prompt 2, the input file consisted of extracted mutation tables from the full document.

### Development and Analysis of General Test Questions

Prompt 1 was developed to query the AI models for correct information about different mutations within the 2023 WHO Mutation Catalogue, containing 30 general test questions. The prompt also specified to keep the inquiry within the supplied document to try avoiding hallucinations. These questions were designed to cover a range of complexity levels, from simple factual queries to more intricate questions requiring interpretation and synthesis of information within the document. For example, we used different questions to probe various aspects of catalogue interpretation, including recalling factual mutation classifications, explaining core concepts (e.g., ‘Uncertain Significance’, Loss of Function (LoF) mutations), retrieving information on specific gene-drug associations, and interpreting complex interactions like epistasis. The questions were also structured to assess the AI models’ ability to handle ambiguous queries and provide accurate information without generating false or misleading content (hallucination). Hallucination was defined as the inclusion of any information presented as ‘factual’ but not found within the provided catalogue document, irrespective of whether the external information was correct. The answers of each model were scored for accuracy, completeness, clarity, citation and hallucinations according to the scale shown in Table 1.

**Table 1.** Evaluation criteria for AI model responses, including descriptions of scoring metrics for accuracy, completeness, clarity, source citation, and avoidance of hallucination.

| **Criterion** | **Description** |
|---|---|
| **Accuracy** | 3 = Fully accurate (completely correct) |
|  | 2 = Mostly accurate (minor inaccuracies) |
|  | 1 = Partially accurate (significant inaccuracies) |
|  | 0 = Inaccurate (completely wrong) |
| **Completeness** | 3 = Fully complete (addresses all aspects) |
|  | 2 = Mostly complete (misses minor details) |
|  | 1 = Partially complete (leaves out significant information) |
|  | 0 = Incomplete (fails to address the question) |
| **Clarity** | 2 = Clear and well-organized (easy to understand) |
|  | 1 = Somewhat clear (could be better organized) |
|  | 0 = Unclear (confusing) |
| **Source Citation** | 2 = Correctly cited (accurate citation) |
|  | 1 = Partially cited (vague or inaccurate citation) |
|  | 0 = Not cited (no citation) |
| **Hallucination** | 0 = No hallucination (only information from the document) |
|  | 1 = Hallucination present (includes information not in the document) |

### Deep Testing of Mutation Search and Retrieval Capabilities

To rigorously evaluate the mutation search and retrieval capabilities of the selected AI models, we employed a two-tiered testing approach:

#### 1. Full catalogue query testing

We compiled a list of 50 distinct mutations from the 2023 WHO Mutation Catalogue. These mutations were chosen to represent a variety of genes, drugs, and levels of association with resistance (e.g., ‘Assoc with R’, ‘Uncertain significance’). Each AI model was queried with Prompt 2, asking it to locate the final confidence grading of each of the 50 individual mutations within the provided catalogue file. For each query, we recorded whether the AI model successfully located and provided accurate information about the mutation. A “success” was recorded if the model correctly interpreted the mutation. A “failure” was recorded if the model could not locate the mutation, provided incorrect information, or hallucinated information not present in the document. The success and failure rates for each AI model across the 50 individual mutation queries were calculated and compared.

#### 2. Antibiotic-specific table query testing

Additionally, as it has been documented that LLM models can get “distracted” by large text files [5], we extracted and prepared a concatenate of the tables for each antibiotic from the 2023 WHO Mutation Catalogue to simplify the input context. This resulted in a considerably smaller file, focused only on the mutations associated with resistance. Similar to the previous approach, we queried the same mutations and recorded successes and failures for each query based on the information retrieved by the AI model. This approach allowed us to assess the AI models’ ability to navigate and get information from smaller, more focused datasets as opposed to the entire document, text and tables.

### Application of Additional Grading Rules to Novel Mutations

To assess the AI models’ ability to apply the 2023 WHO Mutation Catalogue’s additional grading rules[6] beyond the explicitly listed mutations, a third prompt (Prompt 3) was designed. This prompt presented the models with a series of novel mutations not found within the primary catalogue tables. The prompt instructed the models to interpret these mutations based on the grading rules and interpretation criteria detailed in the document. For each novel mutation, the AI models were asked to determine its predicted confidence group, identify the associated drug, and cite the specific rule from the catalogue that justified their interpretation. The answers were again assessed by accuracy, completeness, clarity, citation and hallucinations. This analysis aimed to evaluate the models’ capacity for deductive reasoning and their ability to apply the catalogue’s additional grading rules to previously unseen data.

## Statistical Analysis

Statistical analysis was performed to determine if significant differences existed between the mean performance scores of the four AI models for the metrics evaluated. For Prompt 1 and Prompt 3, one-way Analysis of Variance (ANOVA) tests were conducted separately for each metric (Accuracy, Completeness, Clarity, Source Citation) using the *scipy*.*stats* library in Python. For each test, the individual scores obtained for each question were grouped by AI model. A p-value less than 0.05 was considered statistically significant. ANOVA assumptions, such as homogeneity of variances, were considered, and tests were skipped if assumptions were clearly violated (e.g., zero variance within groups for specific metrics in Prompt 3).

To assess statistical differences in the binary success/failure outcomes of the mutation search and retrieval task (Prompt 2), Pearson’s Chi-squared tests were conducted. These tests compared the proportion of successful retrievals across the four AI models independently for both the full catalogue query testing and the antibiotic-specific table query testing scenarios. Analyses were also performed using the *scipy*.*stats* library in Python, with a p-value threshold of 0.05 for statistical significance.

## RESULTS

This study evaluated the performance of four prominent AI models—Google Gemini 2.5 Pro, OpenAI ChatGPT 4.1, Perplexity AI, and DeepSeek R1—in accurately retrieving and interpreting information from the 2023 WHO Mutation Catalogue for *Mycobacterium tuberculosis* drug resistance. The models were first tested using the standardized Prompt 1 containing 30 questions provided in Supplementary Material 1 and assessed based on five criteria: accuracy, completeness, clarity, source citation, and hallucinations.

### Overall Performance Across AI Models

The performance of each AI model varied across the different evaluation criteria. A summary of the mean scores and standard deviation (SD) for each metric is presented in Table 2.

**Table 2.** Mean ± SD of AI models across the evaluation criteria for Prompt 1 answers.

| **Metric** | **ChatGPT 4.1** | **DeepSeek R1** | **Gemini 2.5 Pro** | **Perplexity AI** |
|---|---|---|---|---|
| **Accuracy (0-3)** | 2.23 ± 1.01 | 2.23 ± 1.06 | 2.77 ± 0.61 | 1.87 ± 1.11 |
| **Completeness (0-3)** | 2.17 ± 1.01 | 2.23 ± 1.04 | 2.80 ± 0.52 | 2.00 ± 1.08 |
| **Clarity (0-2)** | 2.00 ± 0.00 | 1.73 ± 0.53 | 1.87 ± 0.34 | 1.60 ± 0.49 |
| **Source Citation (0-2)** | 0.00 ± 0.00 | 1.77 ± 0.51 | 1.97 ± 0.18 | 1.10 ± 0.64 |
| **Number of Hallucinations** | 0 | 5 | 0 | 8 |

An overall comparison (Figure 1) shows that Google Gemini 2.5 Pro demonstrated the highest accuracy, with a mean score of 2.77 ± 0.61. Then, both ChatGPT 4.1 and DeepSeek R1 performed well with a mean accuracy score of 2.23 ± 1.01 and 2.23 ± 1.06 respectively. Perplexity AI had the lowest accuracy score of 1.87 ± 1.11. Similar to accuracy, Google Gemini 2.5 Pro showed the highest completeness, with a mean score of 2.80 ± 0.52. Again, both ChatGPT 4.1 and DeepSeek R1 followed with completeness scores of 2.17 ± 1.01 and 2.23 ± 1.04 respectively. The clarity of responses was generally high across all models, with ChatGPT 4.1 obtaining a perfect score for this metric. As for citation, Gemini 2.5 Pro and DeepSeek R1 were more effective at citing sources, with mean scores of 1.97 ± 0.18 and 1.77 ± 0.51, respectively. Perplexity AI had a mean citation score of 1.10 ± 0.64, while ChatGPT 4.1 did not cite any precise sources. Finally, both Google Gemini 2.5 Pro and ChatGPT 4.1 tied for the best performance in avoiding hallucination, with a perfect score of zero, indicating no instances of generating false information or information not contained with the 2023 Mutation Catalog. DeepSeek R1 had a total of 5 hallucinations, while Perplexity AI had the highest number, 8 total hallucinations.

**Figure 1.**
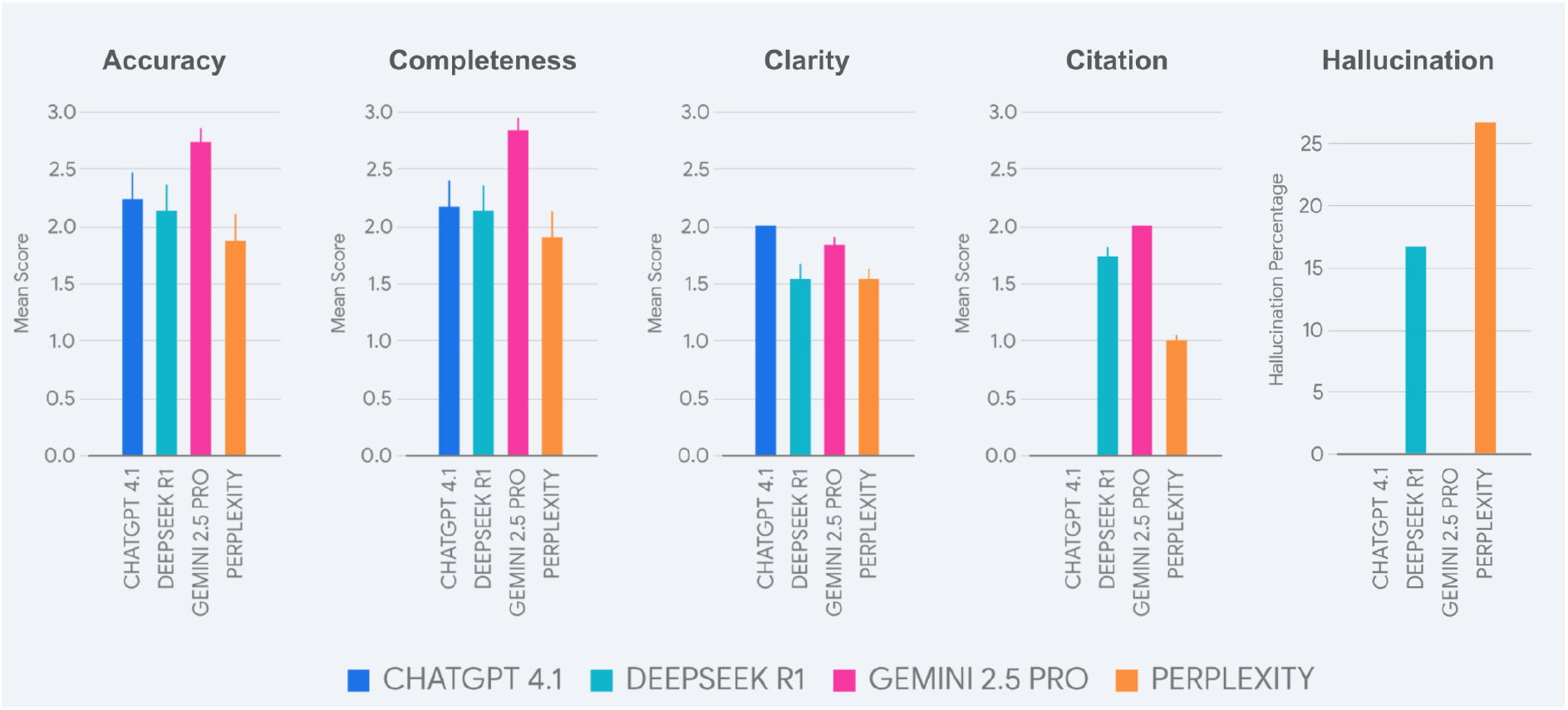
Evaluation of answers to Prompt 1. Mean scores for each metric and AI model except hallucinations, which shows the percentage (total questions = 30). Error bars represent the Standard Error of the Mean (SEM).

To determine if the observed differences in mean scores between the AI models were statistically significant, a one-way ANOVA was performed for each metric evaluated in Prompt 1. The analysis revealed statistically significant differences between the models for all four metrics: accuracy (F(3, 116) = 3.04, p = 0.032), completeness (F(3, 116) = 3.93, p = 0.010), clarity (F(3, 116) = 6.88, p = 0.0003), and source citation (F(3, 116) = 354.39, p < 0.0001). These results confirm that the performance variations noted across the AI models are statistically significant.

### Performance on Specific Questions

The performance of AI models varied significantly across different types of questions. For questions requiring specific factual recall (e.g., “What is the most common resistance-conferring mutation for rifampicin?”), all models performed well. However, for more complex questions requiring interpretation or synthesis of information (e.g., “What is the role of ‘epistasis’ in interpreting resistance to bedaquiline and clofazimine?”), the performance was more variable.

Questions such as “Describe the different groups used to classify mutations in the catalogue” and “Explain what is meant by ‘Uncertain Significance’ mutations” were answered accurately by most models, particularly Gemini 2.5 Pro and DeepSeek R1. Then, questions like “Identify a mutation in the Rv0678 gene that is associated with bedaquiline resistance” and “Locate a mutation that is associated with resistance to linezolid in the rrl gene” showed significant variability in accuracy and completeness across the models. And finally, instances of hallucination were observed across DeepSeek R1 and Perplexity AI, particularly in questions requiring specific gene and mutation information. As stated before, Gemini 2.5 Pro and ChatGPT 4.1 had no hallucinations.

Overall, Google Gemini 2.5 Pro consistently outperformed other models across most evaluation criteria, particularly in accuracy, completeness, and avoidance of hallucination. ChatGPT 4.1 excelled in the clarity of its answers but lacked proper citation functionality. DeepSeek R1 also demonstrated strong performance, especially in accuracy and source citation. Perplexity AI showed variability in their performance, with instances of hallucination and lower accuracy scores compared to the other models. This data provides a quantitative assessment of the AI models’ capabilities in interacting with and interpreting the 2023 WHO Mutation Catalogue, highlighting the strengths and weaknesses of each model in this specific application.

### Search and Retrieval Performance

To rigorously evaluate the mutation search and retrieval capabilities of the four tested AI models, we employed a two-tiered testing approach: full catalogue query testing and antibiotic-specific table query testing.

In the full catalogue query testing, each AI model was tasked with locating the confidence rating of 50 distinct mutations from the entire 2023 WHO Mutation Catalogue. The success and failure rates for each model were recorded. Table 3 summarizes the performance of each AI model in this testing exercise.

**Table 3:** Full catalogue query testing results.

| **AI Model** | **Successes** | **Failures** | **Success Rate (%)** |
|---|---|---|---|
| **Gemini 2.5 Pro** | 39 | 11 | 78 |
| **ChatGPT 4.1** | 25 | 25 | 50 |
| **DeepSeek R1** | 23 | 27 | 46 |
| **Perplexity AI** | 17 | 33 | 34 |

As shown in Table 3, Google Gemini 2.5 Pro demonstrated the highest success rate of 78%, successfully locating information for 39 out of the 50 mutations. ChatGPT 4.1 scored second place with a 50% success rate (25 out of 50). DeepSeek R1 had a 46% success rate (23 out of 50) while Perplexity AI had a 34% success rate (17 out of 50).

In the antibiotic-specific table query testing, separate tables were extracted for each antibiotic from the 2023 WHO Mutation Catalogue. The same 50 mutations were queried within these smaller, drug-specific tables. Table 4 presents the performance of each AI model in this testing exercise.

**Table 4.** Antibiotic-specific table query testing results.

| **AI Model** | **Successes** | **Failures** | **Success Rate (%)** |
|---|---|---|---|
| **DeepSeek R1** | 47 | 3 | 94 |
| **Gemini 2.5 Pro** | 46 | 4 | 92 |
| **ChatGPT 4.1** | 41 | 9 | 82 |
| **Perplexity AI** | 29 | 21 | 58 |

In the antibiotic-specific table query testing, Google Gemini 2.5 Pro and DeepSeek R1 demonstrated very high success rates, with 92% (46 out of 50) and 94% (47 out of 50), respectively. ChatGPT 4.1 achieved a 82% success rate (41 out of 50), and Perplexity AI still had a low 58% success rate (29 out of 50).

Comparing the two testing approaches (Figure 2), it was observed that ChatGPT 4.1, Perplexity AI, and DeepSeek R1 generally performed better in the antibiotic-specific table query testing compared to the full catalogue query testing. This suggests that these models may be more effective at retrieving information from smaller, more focused datasets. Google Gemini 2.5 Pro maintained its high performance across both testing approaches, indicating a robust ability to navigate and retrieve information from both large and small datasets, albeit with a clear advantage when having a smaller data space to search. Chi-squared tests confirmed that the observed differences in success rates for this mutation search and retrieval task were statistically significant. Significant differences between the models’ performance were found both when querying the full catalogue (χ2(3)=20.40, p < 0.001) and when querying the smaller, antibiotic-specific tables (χ2(3)=25.00, p < 0.001). This indicates that the varying abilities of the models to correctly locate mutation information are unlikely due to random chance in either testing scenario.

**Figure 2.**
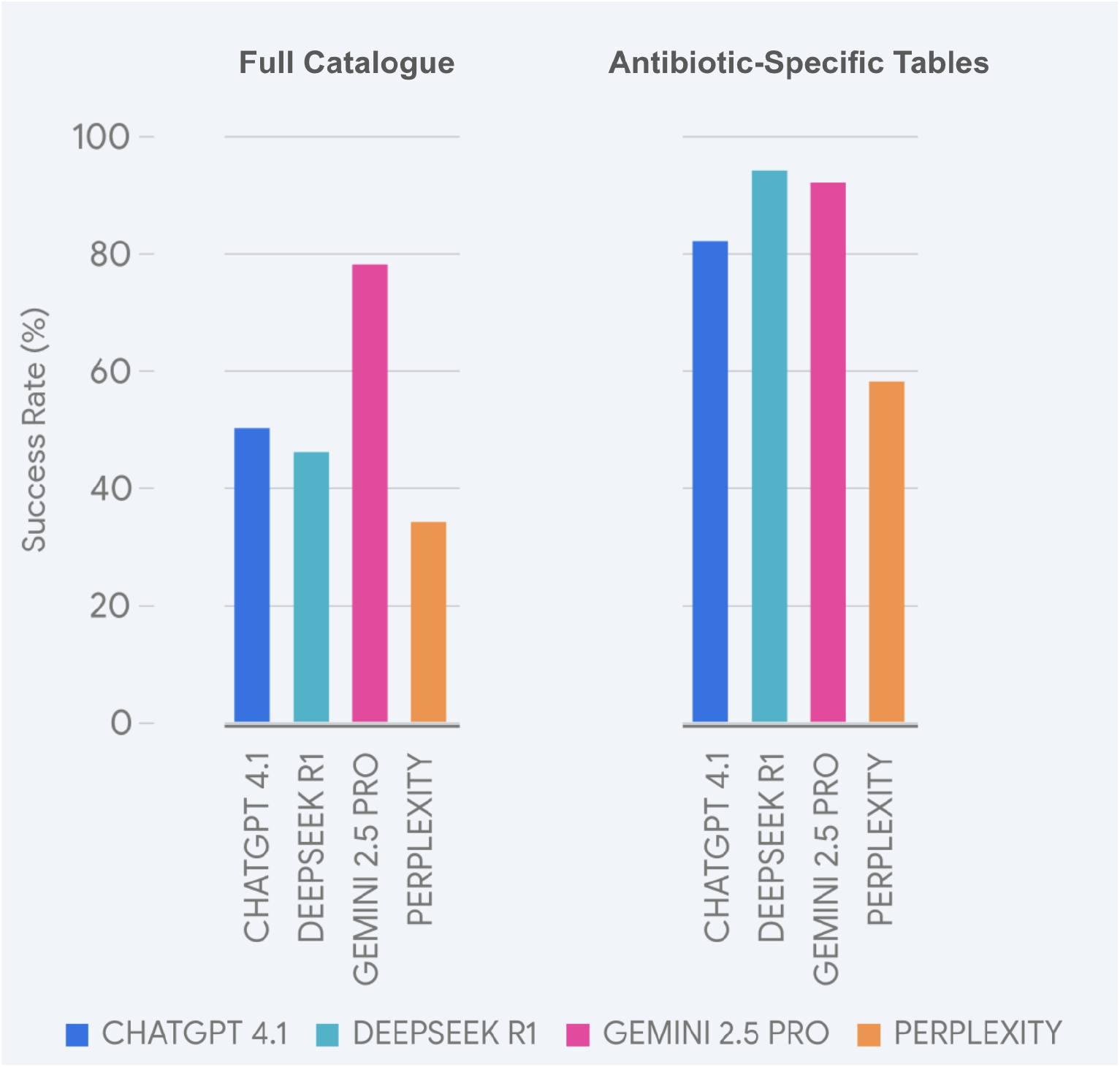
Evaluation of answers to Prompt 2. *Success rates (%) of locating mutations and their confidence ratings in both the full catalogue file and the fragmented antibiotic tables.*

### Additional Grading Rules Evaluation Performance

To evaluate the AI models’ ability to apply the 2023 WHO Mutation Catalogue’s additional grading rules to mutations that are not explicitly listed, Prompt 3 was used, presenting mutations not explicitly contained in the catalogue’s tables for interpretation. This prompt instructed the models to determine the predicted confidence group, identify the associated drugs, and cite the relevant rules. The answers were again evaluated using the scoring criteria from Table 1. A summary of the mean scores and standard deviation (SD) for each metric after this analysis is presented in Table 5.

**Table 5.** Mean ± SD of AI models across the evaluation criteria for Prompt 3 answers.

| **Metric** | **ChatGPT 4.1** | **DeepSeek R1** | **Gemini 2.5 Pro** | **Perplexity AI** |
|---|---|---|---|---|
| **Accuracy (0-3)** | 2.5 ± 1.08 | 2.9 ± 0.32 | 2.4 ± 1.26 | 1.4 ± 0.84 |
| **Completeness (0-3)** | 2.7 ± 0.95 | 2.9 ± 0.32 | 2.7 ± 0.95 | 2.2 ± 0.42 |
| **Clarity (0-2)** | 2.0 ± 0 | 1.9 ± 0.32 | 2.0 ± 0 | 2.0 ± 0 |
| **Source Citation (0-2)** | 1.7 ± 0.67 | 2.0 ± 0 | 1.6 ± 0.84 | 2.0 ± 0 |
| **Number of Hallucinations** | 1 | 0 | 1 | 3 |

The data in Table 5 provides a quantitative comparison of how well each AI model performed in interpreting mutations not explicitly listed in the catalogue, demonstrating their capacity for deductive reasoning and application of the catalogue’s broader guidelines listed as additional grading rules. This overall comparison (Figure 3) shows that DeepSeek R1 demonstrated the highest accuracy, with a mean score of 2.9 ± 0.32. Then, both ChatGPT 4.1 and Google Gemini 2.5 Pro performed well with mean accuracy scores of 2.5 ± 1.08 and 2.4 ± 1.26, respectively. Perplexity AI had the lowest accuracy score of 1.4 ± 0.84. Similar to accuracy, DeepSeek R1 showed the highest completeness, with a mean score of 2.9 ± 0.32. Again, both ChatGPT 4.1 and Google Gemini 2.5 Pro followed with completeness scores of 2.7 ± 0.95. The clarity of responses was generally high across all models, with ChatGPT 4.1 and Google Gemini 2.5 Pro obtaining a perfect score for this metric. As for citation, Google Gemini 2.5 Pro and ChatGPT 4.1 were effective at citing the applied rules, with mean scores of 1.6 ± 0.84 and 1.7 ± 0.67, respectively. DeepSeek R1 and Perplexity AI had a perfect citation score of 2.0 ± 0.00. Finally, DeepSeek R1 had the best performance in avoiding hallucination, with a perfect score of zero, indicating no instances of generating false information. Google Gemini 2.5 Pro and ChatGPT 4.1 had a total of 1 hallucination each, while Perplexity AI had the highest number, 3 total hallucinations.

**Figure 3.**
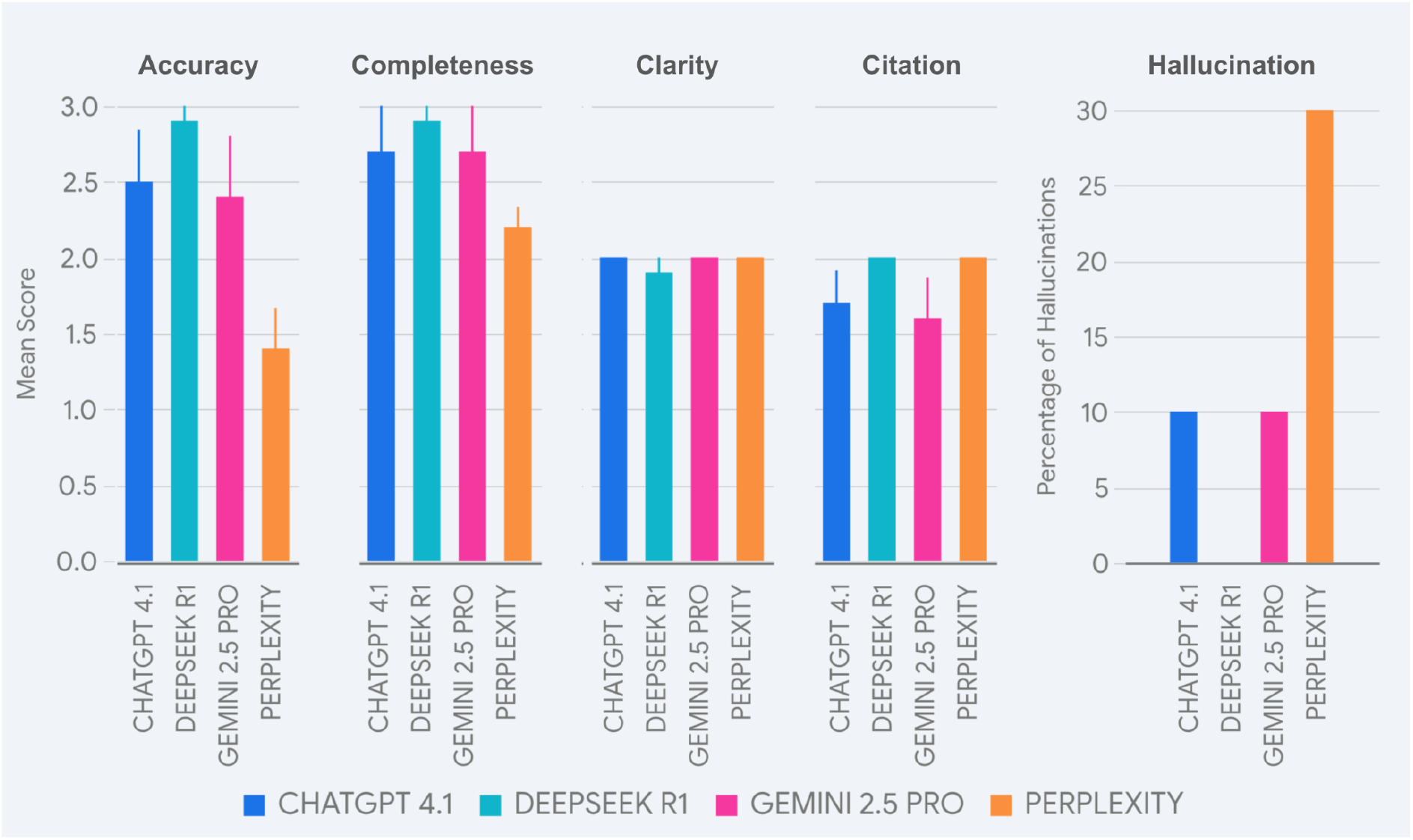
Evaluation of answers to Prompt 3. Mean scores for each metric and AI model except hallucinations, which shows the percentage (total questions = 10). Error bars represent the Standard Error of the Mean (SEM).

Statistical analysis using one-way ANOVA was performed to compare model performance on the additional grading rules task (Prompt 3). A significant difference between models was found for Accuracy (F(3, 36) = 4.55, p = 0.008). However, no significant difference was detected for Completeness (F(3, 36) = 1.72, p = 0.181). ANOVA was not suitable for the Clarity and Source Citation metrics, as the assumption of non-zero variance within groups was violated due to identical scores given by one or more models; thus, p-values could not be calculated for these metrics using this method.

## DISCUSSION

This study aimed to evaluate the potential of prominent AI models (Google Gemini 2.5 Pro, OpenAI ChatGPT 4.1, Perplexity AI, and DeepSeek R1) to facilitate natural language user interactions with the 2023 WHO Mutation Catalogue for *Mycobacterium tuberculosis* drug resistance. Our findings reveal significant variations in the performance of these models across different evaluation criteria and testing approaches.

Google Gemini 2.5 Pro consistently demonstrated superior performance across multiple aspects of the evaluation. It achieved the highest mean scores in accuracy and completeness in the general test questions, indicating its ability to provide correct and comprehensive answers. Additionally, Gemini 2.5 Pro excelled in source citation, demonstrating its skill in backing up its responses with specific references from the document. Crucially, with Prompt 1, no instances of hallucinations were observed in its responses, emphasizing its reliability. In the search and retrieval tests, Gemini 2.5 Pro showed high success rates in both full catalogue queries and antibiotic-specific table queries, suggesting its superior performance in the area of attention [7]. However, it struggled to appropriately apply the additional grading rules in two instances. Overall, this model seemed consistent and accurate, positioning it as one of the strongest candidates for future implementation as a drug resistant-TB mutation interpretation tool. Gemini’s performance may reflect architectural advantages or training data characteristics optimized for processing and synthesizing information from large, dense documents.

OpenAI ChatGPT 4.1 demonstrated notable strengths, particularly in the clarity of its responses, consistently achieving perfect scores in this metric. However, it faced significant limitations in its ability to provide accurate source citations, failing to cite specific sections or pages within the document and only referring to the whole provided document. Additionally, while ChatGPT 4.1 exhibited good accuracy and completeness in general test questions, its performance in mutation search and retrieval tasks was less reliable compared to Google Gemini 2.5 Pro and DeepSeek R1. In the full catalogue query testing, its success rate was only 50%, although it improved to 82% in the antibiotic-specific table query testing. These findings suggest that ChatGPT 4.1, which has been documented to be able to handle up to one million tokens [8], appears to still struggle with larger, less structured datasets. The lack of source citations within the document appears to be a limitation of this model.

DeepSeek R1 demonstrated strong performance, particularly in accuracy and source citation in the general test questions. It also excelled in the antibiotic-specific table query testing, achieving a 94% success rate, indicating its proficiency in navigating focused datasets. Notably, DeepSeek R1 showed significant strength in applying additional grading rules to novel mutations, achieving the highest accuracy in this evaluation, a difference found to be statistically significant compared to other models (p = 0.008), and avoiding hallucinations entirely. However, it exhibited some instances of hallucination in the general test questions, with a total of 5 hallucinations observed, which suggests a need for further refinement to ensure consistent reliability across all tasks. However, DeepSeek’s notable strength in applying grading rules, despite some hallucinations on general queries, suggests potentially stronger capabilities in pattern recognition or logical inference based on defined rules, a valuable trait for interpreting new variants.

Perplexity AI demonstrated the most variable performance among the AI models evaluated in this study. While it showed some proficiency in clarity and source citation in specific instances, it struggled with accuracy and completeness across both general test questions and mutation search/retrieval tasks. Notably, Perplexity AI exhibited the highest number of hallucinations, generating false or misleading information in a significant portion of its responses. Its performance in the full catalogue query testing was particularly low, with a success rate of only 34%. Although its success rate improved in the antibiotic-specific table query testing, it still remained the lowest among the four models. These results suggest that the current Perplexity AI model is not suitable for this study’s use case of interrogating and accessing large, complex technical documents.

This study had several limitations. First, the evaluation was based on a specific set of carefully designed prompts, and it is well documented that the performance of the AI models may vary with different inputs [9]. Second, the study focused on four specific versions of available LLMs, and the results may not be generalizable to all versions of the model. Additionally, given the fast-evolving AI market and products – the performance documented here cannot be extrapolated to newer models from these sources. Third, the evaluation was primarily quantitative, and further qualitative analysis of the AI responses could provide deeper insights into their strengths and weaknesses.

Nevertheless, the results from evaluating these AI models on their ability to interact with the 2023 WHO Mutation Catalogue reveal significant implications for developing a custom AI agent for this use case. Notably, Google Gemini 2.5 Pro demonstrated superior performance in accuracy, completeness, and avoidance of hallucinations across general test questions and search/retrieval tasks, making it a strong candidate for the core engine of such an agent. Its ability to effectively navigate both full catalogue queries and antibiotic-specific table queries, along with its high rate of successful source citation, indicates a robust understanding of the complex data structure and strong fidelity to the material endorsed in the Mutation Catalogue. It is plausible that with additional training on more fragmented components of the Mutation Catalogue, that this tool could reach near perfect fidelity and accuracy – potentially sufficient for use as a clinical decision support tool.

ChatGPT 4.1 also presented itself as a good candidate, although its poor scores on data retrieval pose serious limitations unless these can also be overcome with more fragmented data inputs. DeepSeek R1 also showed promise, particularly in applying additional grading rules to novel mutations and achieving high accuracy in focused datasets, suggesting it could enhance the agent’s deductive reasoning capabilities. However, the performance variability and higher hallucination rates observed with Perplexity AI suggest this use case is currently not within its capabilities.

The overall improvement of search and retrieval performance across all models when using the antibiotic-specific table queries, instead of the whole catalogue, implies that structuring the catalogue into smaller, more manageable datasets could significantly improve information retrieval efficiency for any future agent and the need for model improvement in this area. Given the significant negative potential of hallucination risk, only agents demonstrated to have fidelity to the endorsed material (in this case the 2023 Mutation Catalogue) should be considered appropriate for this use case and emphasize the need for internal document citation capabilities as an ongoing quality control measure. Ultimately, a custom AI agent built on these insights could greatly enhance the accessibility and utility of the WHO Mutation Catalogue. An agent optimized for this purpose could provide a broader audience with an intuitive, natural multi-language tool for efficient information retrieval, nuanced mutation interpretation, and personalized decision support for clinical interpretation of complex mutations from sequencing and other molecular diagnostics, especially coupled with the WHO treatment guidelines for TB [10, 11].

Future research should explore the use of AI models for more advanced tasks, such as automatically classifying mutations from sequencing data or generating personalized reports based on user queries [12]. It would be valuable to investigate how AI tools, trained on highly curated knowledgebases can be integrated directly into existing bioinformatic workflows for use with antimicrobial resistance (AMR) and oncology diagnostics towards precision medicine.

In conclusion, this study demonstrates the potential of existing LLMs to facilitate non-expert, natural language user interaction with the 2023 WHO Mutation Catalogue for drug resistant TB. The findings underscore the importance of careful model selection and rigorous testing when deploying AI tools for scientific information retrieval and interpretation. As AI technology continues to advance, it has the potential to significantly enhance the accessibility and utility of critical, complex and highly curated resources like the WHO Mutation Catalogue, ultimately contributing to improved TB management and care.

## SUPPLEMENTARY MATERIAL

### Prompt 1

“You are an expert in interpreting the WHO Mutation Catalogue for *Mycobacterium tuberculosis* drug resistance. I am providing you with the document ‘catalogue_2023.pdf’. Your task is to accurately answer the following questions based solely on the information contained within this document. Please provide detailed and specific answers, citing the relevant sections or pages where you found the information whenever possible. Avoid making assumptions or providing information that is not explicitly stated in the document. If a question cannot be answered using the document, please state ‘Information not found in document.’ Here are the questions:

1. Describe the different groups used to classify mutations in the catalogue.
2. How are mutations graded in the catalogue?
3. Explain what is meant by “Uncertain Significance” mutations.
4. How do you interpret a mutation listed as “Assoc w R-interim”?
5. Explain the concept of “loss-of-function” (LoF) mutations and their relevance to drug resistance.
6. Does the catalogue contain mutations that are not associated with drug resistance?
7. What is the most common *resistance-conferring mutation for rifampicin?*
8. *Identify the nucleotide change associated with the most frequent inhA promoter mutation linked to isoniazid resistance*
9. *What is the role of “epistasis” in interpreting resistance to bedaquiline and clofazimine?*
10. *Identify a mutation in the Rv0678 gene that is associated with bedaquiline resistance*.
11. *Are insertions or deletions (indels) in the pncA gene known to cause pyrazinamide resistance?*
12. *Locate a mutation that is associated with resistance to linezolid in the rrl gene*.
13. *What is the codon position of the most common pyrazinamide resistance mutation in the pncA gene?*
14. *How does the tlyA gene contribute to capreomycin resistance?*
15. *Provide an example of a compensatory mutation in the rpoC gene that is found in rifampicin-resistant strains*.
16. *What genes are involved in bedaquiline resistance?*
17. *A Trp328Leu mutation in katG has been found in a sample. What does it mean?*
18. *Explain the potential impact of a mutation at codon 94 of gyrA*.
19. *Does the Leu116Pro mutation in pncA confer resistance to pyrazinamide?*
20. *What is the significance of the embB M306V mutation in ethambutol resistance?*
21. How does the rpoB S450L mutation lead to rifampicin resistance at the molecular level?
22. Find an example of a mutation in the *rpsL* gene that is associated with streptomycin resistance.
23. Is the embA regulatory region involved in ethambutol resistance?
24. Are there any mutations outside the rifampicin resistance-determining region (RRDR) of rpoB that contribute to resistance?
25. Find an example of a mutation in the pepQ gene that has been linked to resistance against delamanid.
26. Identify a mutation in the *mmpL5* gene that interacts with *Rv0678* mutations in the context of bedaquiline resistance.
27. Does the pncA gene contain any known duplications that contribute to pyrazinamide resistance?
28. Compare the impact of the katG S315T versus an inhA −15C>T on isoniazid resistance.
29. How does a frameshift mutation in pncA compare to a missense SNP in pncA in terms of pyrazinamide resistance?
30. Are there any documented instances of large deletions in the katG gene, and do they impact isoniazid resistance?”

### Prompt 2

#### Full catalogue search

You are an expert in interpreting the WHO Mutation Catalogue for Mycobacterium tuberculosis drug resistance. I am providing you with the document ‘catalogue_2023.pdf’. Your task is to locate the specific confidence grading (e.g., ‘Assoc w R’, ‘Uncertain Significance’, ‘Not assoc w R’, etc.) for each of the mutations listed below, based solely on the information contained within the provided document. Do not infer grading based on rules or other mutations; only report what is explicitly stated for the listed mutations. Avoid making assumptions or providing information not present in the document.

Here is the list of mutations: ‘rpoB_p.His445Leu’, ‘rpoB_p.Val170Phe’, ‘rpoB_p.Met434Val’, ‘rpoC_p.Glu1092Asp’, ‘katG_p.Ser315Thr’, ‘katG_LoF’, ‘inhA_c.-777C>T’, ‘inhA_c.-154G>A’, ‘ahpC_c.-88G>A’, ‘rrs_n.1402C>T’, ‘embB_p.Met306Val’, ‘embB_p.Leu74Arg’, ‘embA_c.-12C>T’, ‘embC_c.-1188C>T’, ‘ubiA_p.Val49lle’, ‘pncA_LoF’, ‘pncA_p.His57Asp’, ‘pncA_p.Val139Ala’, ‘pncA_p.Ile133Thr’, ‘pncA_c.138A>G’, ‘rpsA_p.Met432Thr’, ‘gyrA_p.Asp94Gly’, ‘gyrA_p.Ala90Val’, ‘gyrB_p.Asp461Asn’, ‘gyrB_p.Asn499Thr’, ‘gyrA_p.Ser95Thr’, ‘Rv0678_LoF’, ‘Rv0678_p.Gly121Arg’, ‘pepQ_LoF’, ‘atpE_p.Ala63Pro’, ‘mmpL5_LoF’, ‘rplC_p.Cys154Arg’, ‘rrl_n.2814G>T’, ‘rrl_n.2270G>T’, ‘tsnR_p.Leu232Pro’, ‘Rv1979c_c.-129A>G’, ‘ddn_LoF’, ‘ddn_p.Leu49Pro’, ‘fbiC_LoF’, ‘fbiA_LoF’, ‘fbiA_p.Arg321Ser’, ‘Rv2983_LoF’, ‘rrs_n.1401A>G’, ‘rrs_n.1484G>T’, ‘eis_c.-14C>T’, ‘rpsL_p.Lys43Arg’, ‘gid_p.Glu99* ‘, ‘gid_LoF’, ‘ethA_LoF’, ‘mshA_p.Ala187Val’.

#### Antibiotic-specific tables

You are an expert in interpreting the WHO Mutation Catalogue for Mycobacterium tuberculosis drug resistance. I am providing you with an extract from the catalogue containing only the tables with resistance mutations for each antibiotic. Your task is to locate the specific confidence grading (e.g., ‘Assoc w R’, ‘Uncertain Significance’, ‘Not assoc w R’, etc.) for each of the mutations listed below, based solely on the information contained within the provided document. Do not infer grading based on rules or other mutations; only report what is explicitly stated for the listed mutations. Avoid making assumptions or providing information not present in the document.

Here is the list of mutations: ‘rpoB_p.His445Leu’, ‘rpoB_p.Val170Phe’, ‘rpoB_p.Met434Val’, ‘rpoC_p.Glu1092Asp’, ‘katG_p.Ser315Thr’, ‘katG_LoF’, ‘inhA_c.-777C>T’, ‘inhA_c.-154G>A’, ‘ahpC_c.-88G>A’, ‘rrs_n.1402C>T’, ‘embB_p.Met306Val’, ‘embB_p.Leu74Arg’, ‘embA_c.-12C>T’, ‘embC_c.-1188C>T’, ‘ubiA_p.Val49lle’, ‘pncA_LoF’, ‘pncA_p.His57Asp’, ‘pncA_p.Val139Ala’, ‘pncA_p.Ile133Thr’, ‘pncA_c.138A>G’, ‘rpsA_p.Met432Thr’, ‘gyrA_p.Asp94Gly’, ‘gyrA_p.Ala90Val’, ‘gyrB_p.Asp461Asn’, ‘gyrB_p.Asn499Thr’, ‘gyrA_p.Ser95Thr’, ‘Rv0678_LoF’, ‘Rv0678_p.Gly121Arg’, ‘pepQ_LoF’, ‘atpE_p.Ala63Pro’, ‘mmpL5_LoF’, ‘rplC_p.Cys154Arg’, ‘rrl_n.2814G>T’, ‘rrl_n.2270G>T’, ‘tsnR_p.Leu232Pro’, ‘Rv1979c_c.-129A>G’, ‘ddn_LoF’, ‘ddn_p.Leu49Pro’, ‘fbiC_LoF’, ‘fbiA_LoF’, ‘fbiA_p.Arg321Ser’, ‘Rv2983_LoF’, ‘rrs_n.1401A>G’, ‘rrs_n.1484G>T’, ‘eis_c.-14C>T’, ‘rpsL_p.Lys43Arg’, ‘gid_p.Glu99* ‘, ‘gid_LoF’, ‘ethA_LoF’, ‘mshA_p.Ala187Val’.

### Prompt 3

“You are an expert in interpreting the WHO Mutation Catalogue for *Mycobacterium tuberculosis* drug resistance. I am providing you with the document ‘catalogue_2023.pdf’. Your task is to accurately answer the following questions based solely on the information contained within this document. You may apply the additional grading rules and other interpretation criteria detailed in the document (particularly Section 2.1, Table 1) for mutations which are not found in the main catalogue tables. For each mutation, state its predicted confidence group (1-5), the associated drug(s) and the specific rule(s) from the catalogue that justify your interpretation. Here are the questions:

1. A novel frameshift mutation, katG_p.Lys154fs, has been detected. How should this be interpreted for isoniazid resistance?
2. A nonsense mutation, pncA_p.Arg140Ala, has been identified in a clinical isolate. What is the expected impact on pyrazinamide susceptibility?
3. A sequencing report has identified a start-loss mutation, Rv0678_p.Met1?, in an isolate. How can this mutation affect the drug resistance profile of the isolate?
4. An isolate contains a large deletion resulting in the loss of the entire fbiC gene. What is the predicted effect on delamanid resistance?
5. A non-synonymous mutation, rpoB_p.Gln432Arg, located within the Rifampicin Resistance-Determining Region (RRDR), has been found. How should this be classified for rifampicin resistance?
6. The promoter mutation fabG1_c.-8T>A has been detected. What are the implications for isoniazid and ethionamide resistance?
7. The mutation gyrA_p.Gly88Ala has been found in an isolate. How would this be interpreted regarding resistance to fluoroquinolones?
8. An isolate has both the eis_c.-14C>T promoter mutation and an eis_p.Trp50fs frameshift mutation. Will it be resistant to amikacin?
9. A novel synonymous mutation resulting in gyrB_p.Gly520Gly has been detected. How would this mutation affect resistance to fluoroquinolones?
10. An isolate has both the Rv0678_p.Gly121Arg mutation and an mmpL5_LoF mutation. Will it be resistant to bedaquiline?”

### Model Responses

#### GEMINI - PROMPT 1

Based on the information found in the ‘Catalogue of mutations in Mycobacterium tuberculosis complex and their association with drug resistance, second edition’ (catalogue_2023.pdf), here are the answers to your questions:

#### Mutation Classification and Interpretation

- **Classification Groups:** Mutations in the catalogue are classified into five groups based on the statistical evidence supporting their association with phenotypic resistance:
  ∘ **Group 1: Associated with resistance (Assoc w R)** - Strong evidence linking the mutation to resistance. Interpreted as markers of resistance.
  ∘ **Group 2: Associated with resistance-interim (Assoc w R-interim)** - Interim association with resistance, often based on relaxed criteria, WHO-endorsed assay detection, cross-resistance rules, or selection experiments. Also interpreted as markers of resistance.
  ∘ **Group 3: Uncertain significance** - Insufficient or conflicting evidence to determine association with resistance.
  ∘ **Group 4: Not associated with resistance-interim (Not assoc w R-interim)** - Interim evidence suggesting no association with resistance. Not interpreted as markers of resistance.
  ∘ **Group 5: Not associated with resistance (Not assoc w R)** - Strong evidence indicating no association with resistance. Not interpreted as markers of resistance.
- **Mutation Grading:** Mutations are graded into one of the five groups based on statistical analysis of genotype-phenotype data. Key criteria include the number of isolates with the mutation as the sole resistance marker (SOLO), the Positive Predictive Value (PPV) of the SOLO mutation, the Odds Ratio (OR) for resistance, and statistical significance. Additional rules (e.g., based on LoF, WHO precedents, assay detection, cross-resistance) are also applied to finalize the grading.
- **“Uncertain Significance” Mutations:** These are Group 3 mutations. The available evidence from the analysed dataset is insufficient or contradictory to confidently classify them as either associated or not associated with phenotypic drug resistance.
- **“Assoc w R-interim” Interpretation:** A mutation listed as “Assoc w R-interim” (Group 2) should be interpreted as a marker of clinically relevant phenotypic resistance. This grading indicates an association with resistance, but the evidence might stem from sources other than meeting the strict statistical criteria for Group 1, such as detection by WHO-endorsed assays, application of cross-resistance rules, evidence from selection experiments, or meeting relaxed statistical criteria (e.g., for *pncA*).
- **Loss-of-Function (LoF) Mutations:** LoF mutations are genetic changes predicted to cause a complete loss of function of the encoded protein. In this catalogue, this typically includes frameshift mutations, premature stop codons (nonsense), start codon loss mutations, and whole gene deletions (‘feature_ablation’). Inframe insertions/deletions were excluded from the LoF definition in this edition. LoF mutations in specific genes known to be involved in drug activation (e.g., *katG* for INH, *pncA* for PZA, *ethA* for ETO) or drug target modification/efflux regulation (e.g., *gid* for STM, *tlyA* for CAP, *Rv0678* and *pepQ* for BDQ/CFZ, prodrug activators like *ddn* for DLM/PMD) are often directly associated with resistance via additional grading rules.
- **Mutations Not Associated with Resistance:** Yes, the catalogue contains mutations classified in Group 4 (Not assoc w R-interim) and Group 5 (Not assoc w R). These are not considered markers of clinical resistance. Group 4 often includes silent (synonymous) mutations not meeting Group 5 criteria. Group 5 includes mutations identified as neutral based on specific criteria or literature evidence.

#### Specific Drug Resistance Mutations

- **Most Common Rifampicin (RIF) Mutation:** The *rpoB* p.Ser450Leu mutation was the most common, found in 64.4% of RIF-resistant isolates in this dataset.
- **Most Frequent** *inhA* **Promoter Mutation (Isoniazid - INH):** The c.−15C>T mutation (also known as fabG1 c.-15C>T) in the promoter region upstream of *inhA* is the most frequently listed *inhA* promoter mutation associated with INH resistance in the provided table.
- **Role of “Epistasis” (Bedaquiline - BDQ / Clofazimine - CFZ):** Epistasis occurs when the functional consequence of a mutation in one gene is dependent on the genetic background, specifically the presence of mutations in another gene. For BDQ and CFZ, mutations in the repressor gene *Rv0678* typically cause resistance by upregulating the MmpS5/MmpL5 efflux pump. However, if a loss-of-function (LoF) mutation occurs in the *mmpL5* gene (encoding part of the pump) alongside an *Rv0678* mutation, the pump is non-functional, and the *Rv0678* mutation cannot confer resistance.
- *Rv0678* **Mutation (BDQ Resistance):** Examples listed include p.Glu49fs, p.Asp47fs, p.Ile67fs, p.Gly121Arg, p.Leu117Arg, p.Met146Thr, p.Ile67Ser, p.Cys46Arg, p.Ala36Val, p.Asn70Asp, p.Leu32Ser. Many others causing LoF are grouped under Rv0678_LoF.
- *pncA* **Indels (Pyrazinamide - PZA Resistance):** Yes, insertions or deletions (indels) in *pncA* can cause PZA resistance. Frameshift indels are typically considered LoF mutations and fall under the Group 2 LoF rule for *pncA*. However, inframe indels are less likely to confer resistance; for example, pncA p.Gly113_Leu116delinsVal is classified as Group 4 (Not assoc w R-interim).
- *rrl* **Mutation (Linezolid - LZD Resistance):** Examples associated with LZD resistance include mutations at nucleotide positions n.2814G>T, n.2270G>T, n.2269_2270insT, and n.2299G>T. Additional mutations (n.2270G>C, n.2689A>T, n.2746G>A) were included based on selection experiments.
- **Most Common** *pncA* **Codon Position (PZA Resistance):** Mutations at codon 57 (e.g., p.His57Asp, p.His57Arg) are frequently listed as Group 1 resistance mutations. The p.His57Asp mutation is characteristic of intrinsically resistant *M. bovis*. The document does not explicitly state a single most common codon overall for acquired resistance.
- *tlyA* **Role (Capreomycin - CAP Resistance):** Loss-of-function (LoF) mutations in *tlyA* are associated with CAP resistance and classified as Group 2 based on the LoF additional grading rule.
- **Compensatory Mutation (***rpoC* **for RIF Resistance):** Information not found in document. *rpoC* is listed as a Tier 2 gene for RIF resistance, but specific compensatory mutations within it are not provided as examples.
- **Genes Involved in BDQ Resistance:** Tier 1 genes include *Rv0678, atpE*, and *pepQ*. Tier 2 genes include *mmpL5* and *mmpS5. Rv1979c* is also mentioned in relation to BDQ/CFZ.
- *katG* **Trp328Leu Mutation:** The *katG* p.Trp328Leu mutation is classified as Group 1 (Assoc w R) for INH resistance and is assumed to confer high-level resistance.
- **Potential Impact of** *gyrA* **Codon 94 Mutation:** Mutations at *gyrA* codon 94 (Asp94) are associated with resistance to fluoroquinolones (LFX and MFX). Specific mutations like p.Asp94Asn, p.Asp94Gly, p.Asp94His, and p.Asp94Tyr are assumed to confer high-level MFX resistance, while others like p.Asp94Ala confer low-level MFX resistance. Multiple linked low-level mutations can confer high-level resistance.
- *pncA* **Leu116Pro Mutation:** Yes, the *pncA* p.Leu116Pro mutation is classified as Group 2 (Assoc w R-interim) and confers resistance to PZA.
- **Significance of** *embB* **M306V (Ethambutol - EMB Resistance):** The *embB* p.Met306Val mutation is a Group 1 (Assoc w R) mutation for EMB resistance. It is frequently observed and known to confer MICs near the critical concentration, which can sometimes lead to discordance with phenotypic DST results.
- *rpoB* **S450L Molecular Mechanism:** Information not found in document. While identified as the most common RIF resistance mutation, the detailed molecular interaction with the drug is not described.
- *rpsL* **Mutation (Streptomycin - STM Resistance):** Examples include p.Lys43Arg and p.Lys88Arg, both classified as Group 1 (Assoc w R).
- *embA* **Regulatory Region (EMB Resistance):** Yes, the *embA* upstream region is implicated. The mutation embA c.-12C>T, previously Group 1, is now classified as Group 3 (Uncertain significance).
- **Mutations Outside RRDR (***rpoB***) for RIF Resistance:** Yes, *rpoB* p.Val170Phe and p.Ile491Phe are classified as Group 1 (Assoc w R) despite being outside the RRDR (codons 426-452).
- *pepQ* **Mutation (Delamanid - DLM Resistance):** Information not found in document. LoF mutations in *pepQ* are associated with BDQ/CFZ resistance, but no specific link to DLM resistance is mentioned in the provided tables.
- *mmpL5* **Mutation Interacting with** *Rv0678* **(BDQ Resistance):** Loss-of-function (LoF) mutations in *mmpL5* (grouped as ‘mmpL5_LoF’) interact epistatically with *Rv0678* mutations, abrogating the resistance conferred by *Rv0678* mutations if they are genetically linked.
- *pncA* **Duplications (PZA Resistance):** Information not found in document. The document focuses on point mutations, indels, and LoF mutations within *pncA*.
- **Impact Comparison:** *katG* **S315T vs**. *inhA* **-15C>T (INH Resistance):** The *katG* p.Ser315Thr mutation is assumed to confer high-level INH resistance. The *inhA* c.-15C>T mutation (also known as fabG1 c.-15C>T) is assumed to confer low-level INH resistance if it occurs alone.
- **Impact Comparison: Frameshift vs. Missense SNP in** *pncA* **(PZA Resistance):** Frameshift mutations in *pncA* typically lead to a loss of function (LoF) and are associated with PZA resistance (usually Group 2 via the LoF rule) ^1^. Missense Single Nucleotide Polymorphisms (SNPs) result in an amino acid substitution, the impact of which varies greatly depending on the specific change and its location within the PncA enzyme; some missense mutations strongly confer resistance (Group 1 or 2), while others have uncertain significance (Group 3) or are not associated with resistance (Group 4 or 5) ^2^. Generally, LoF mutations like frameshifts are more reliably predicted to cause resistance than missense mutations, whose effects are more variable.
- **Large Deletions in** *katG* **(INH Resistance):** Yes, large deletions affecting *katG* are documented. The bioinformatics pipeline was improved to handle large deletions, and gene deletions (‘feature_ablation’) are considered LoF mutations. LoF mutations in *katG* are associated with (usually high-level) INH resistance via an additional grading rule.

### GEMINI - PROMPT 3

Based on the WHO Mutation Catalogue 2023 provided (‘catalogue_2023.pdf’), here is the interpretation for each mutation:

#### 1. katG_p.Lys154fs (Isoniazid resistance)

∘ **Confidence Group:** 2 (Associated with resistance-interim)
∘ **Drug(s):** Isoniazid (INH)
∘ **Rule(s):** This is a frameshift mutation, which is considered a Loss-of-Function (LoF) mutation. The drug-specific “additional grading rule” states that LoF mutations in *katG* are classified in Group 2. The “other interpretation criteria” specify that *katG* mutations are assumed to confer high-level INH resistance.

#### 2. pncA_p.Arg140Ala (Pyrazinamide susceptibility)

∘ **Confidence Group:** 3 (Uncertain significance)
∘ **Drug(s):** Pyrazinamide (PZA)
∘ **Rule(s):** This specific missense mutation is listed in Table 9 (Abridged variant classification for PZA) with a final confidence grading of Group 3. No additional grading rules apply to change this classification.

#### 3. Rv0678_p.Met1? (Drug resistance profile)

∘ **Confidence Group:** 2 (Associated with resistance-interim)
∘ **Drug(s):** Bedaquiline (BDQ) and Clofazimine (CFZ)
∘ **Rule(s):** This is a start-loss mutation, considered a Loss-of-Function (LoF) mutation. The drug-specific “additional grading rule” classifies LoF mutations in *Rv0678* into Group 2 for BDQ and CFZ resistance. Rv0678 mutations confer cross-resistance to BDQ and CFZ. **Important Caveat:** The “other interpretation criteria” include an epistasis rule stating that *Rv0678* mutations cannot confer resistance if genetically linked with LoF variants in *mmpL5*. Therefore, the impact on the drug resistance profile depends on the status of the *mmpL5* gene.

#### 4. Large deletion of entire fbiC gene (Delamanid resistance)

∘ **Confidence Group:** 2 (Associated with resistance-interim)
∘ **Drug(s):** Delamanid (DLM) and Pretomanid (PMD)
∘ **Rule(s):** Deletion of an entire gene is a Loss-of-Function (LoF) mutation. The drug-specific “additional grading rule” classifies LoF mutations in *fbiC* into Group 2 for DLM and PMD resistance.

#### 5. rpoB_p.Gln432Arg (Rifampicin resistance)

∘ **Confidence Group:** 2 (Associated with resistance-interim)
∘ **Drug(s):** Rifampicin (RIF) and Rifapentine (RPT)
∘ **Rule(s):** This mutation is located within the Rifampicin Resistance-Determining Region (RRDR, codons 426-452) of *rpoB*. The drug-specific “additional grading rule” classifies any non-silent variant in the RRDR of *rpoB* into Group 2, assuming resistance in the absence of contrary evidence. The classification for RIF also applies to RPT.

#### 6. fabG1_c.-8T>A (Isoniazid and Ethionamide resistance)

∘ **Confidence Group:** 1 (Associated with resistance)
∘ **Drug(s):** Isoniazid (INH) and Ethionamide (ETO)/Prothionamide (PTO)
∘ **Rule(s):** This specific promoter mutation (alias *inhA_c*.*-777C>T*) is listed as Group 1 in Table 7 (INH) and Table 19 (ETO). The “other interpretation criteria” state that *fabG1-inhA* mutations confer cross-resistance to ETO and INH and are assumed to confer low-level INH resistance. The ETO classification also applies to PTO.

#### 7. gyrA_p.Gly88Ala (Fluoroquinolone resistance)

∘ **Confidence Group:** 2 (Associated with resistance-interim)
∘ **Drug(s):** Levofloxacin (LFX) and Moxifloxacin (MFX)
∘ **Rule(s):** This mutation is listed as Group 2 in Table 10 (LFX) and Table 11 (MFX) based on additional grading rules (Previous WHO guidance or FQ cross-resistance). *gyrA* mutations confer cross-resistance to LFX and MFX. According to “other interpretation criteria”, this mutation is assumed to confer low-level MFX resistance.

#### 8. eis_c.-14C>T AND eis_p.Trp50fs (Amikacin resistance)

∘ **Confidence Group:** The combination suggests **no resistance** conferred by these mutations due to epistasis. *eis_c*.*-14C>T* alone is Group 2; *eis_p*.*Trp50fs* is an LoF mutation, which does not confer AMK resistance.
∘ **Drug(s):** Amikacin (AMK)
∘ **Rule(s):** The drug-specific “other interpretation criteria” include an epistasis rule: *eis* promoter mutations (like c.-14C>T) cannot confer resistance if genetically linked with LoF variants in the *eis* coding region (like p.Trp50fs). The LoF mutation abrogates the resistance effect of the promoter mutation.

#### 9. gyrB_p.Gly520Gly (Fluoroquinolone resistance)

∘ **Confidence Group:** 4 (Not associated with resistance-interim)
∘ **Drug(s):** Levofloxacin (LFX), Moxifloxacin (MFX)
∘ **Rule(s):** This is a synonymous (silent) mutation. The general “additional grading rule” states that any novel silent variant is classified in Group 4.

#### 10. Rv0678_p.Gly121Arg AND mmpL5_LoF (Bedaquiline resistance)

∘ **Confidence Group:** The combination suggests **no resistance** conferred by these mutations due to epistasis. *Rv0678_p*.*Gly121Arg* alone is Group 1; *mmpL5_LoF* alone is Group 3 but known to cause epistasis.
∘ **Drug(s):** Bedaquiline (BDQ)

**Rule(s):** The drug-specific “other interpretation criteria” include an epistasis rule: *Rv0678* mutations (like p.Gly121Arg) cannot confer resistance if genetically linked with LoF variants in *mmpL5*. The *mmpL5* LoF mutation abrogates the effect of the *Rv0678* resistance mutation.

## DEEPSEEK - PROMPT 1

### Answers Based on the WHO Mutation Catalogue (2023)

#### 1. Different Groups for Mutation Classification

Mutations are stratified into five groups:

- **Group 1 (Assoc w R)**: Strong evidence of association with resistance.
- **Group 2 (Assoc w R-interim)**: Interim association, often based on LoF rules or preliminary data.
- **Group 3 (Uncertain significance)**: Insufficient evidence for classification.
- **Group 4 (Not assoc w R-interim)**: Interim evidence against association.
- **Group 5 (Not assoc w R)**: Confirmed non-resistance markers. *(Pages 15–17, Section 2*.*1; Table 3, Page 23)*

#### 2. Mutation Grading Criteria

Grading relies on statistical thresholds (e.g., PPV *≥* 25%, OR > 1) and validation via WHO-endorsed assays or literature. Groups 1/2 require *≥*5 isolates with SOLO mutations. LoF rules and cross-resistance criteria are applied. *(Section 5*.*7–5*.*8, Pages 97–101)*

#### 3. “Uncertain Significance” Mutations

Group 3 mutations lack sufficient phenotypic or statistical evidence to confirm or refute resistance. Further research is needed. *(Page 16, Section 2*.*1; Table 3, Page 23)*

#### 4. Interpreting “Assoc w R-interim”

Group 2 mutations have interim evidence (e.g., LoF in *katG* for INH or *pncA* for PZA) but require additional validation. They are treated as resistance markers until further data. *(Table 1, Page 16; Section 3*.*3, Page 26)*

#### 5. Loss-of-Function (LoF) Mutations

LoF mutations disrupt gene function (e.g., frameshifts, premature stop codons) and are linked to resistance for prodrugs (e.g., *pncA* for PZA, *ddn* for delamanid). Inframe indels are excluded from LoF classification. *(Section 3*.*4, Page 33; Section 5*.*6, Page 87)*

#### 6. Non-Resistance-Associated Mutations

Yes. Groups 4 and 5 list mutations not associated with resistance (e.g., silent variants, neutral mutations). Example: *rpoB* Glu250Gly (Group 5). *(Table 3, Page 23; Section 2*.*1)*

#### 7. Most Common Rifampicin Resistance Mutation

**rpoB Ser450Leu** (64.4% sensitivity, 99.3% specificity). *(Table 6, Page 30; Section 3*.*2, Page 29)*

#### 8. inhA Promoter Mutation for INH Resistance

**c.-15C>T** (inhA promoter) is a frequent mutation causing low-level INH resistance. *(Table 7, Page 37; Section 3*.*3, Page 26)*

#### 9. Epistasis in Bedaquiline/Clofazimine Resistance

Mutations in *Rv0678* (efflux pump regulator) confer resistance unless accompanied by LoF in *mmpL5* (epistasis). *mmpL5* LoF negates *Rv0678*-mediated resistance. *(Section 3*.*7, Page 52; Table 1, Page 16)*

#### 10. Rv0678 Mutation Example

**Rv0678 frameshift or truncation mutations** (e.g., Gly20*) are Group 2 markers for bedaquiline resistance. *(Section 3*.*7, Page 52; Table 1, Page 16)*

#### 11. Indels in pncA and PZA Resistance

Yes. LoF indels in *pncA* (e.g., frameshifts) are classified as Group 1/2 for PZA resistance. *(Section 3*.*5, Page 36; Table 3, Page 23)*

#### 12. Linezolid Resistance Mutation in rrl

**rrl C154R** (23S rRNA) is a Group 1 mutation associated with linezolid resistance. *(Section 3*.*8, Page 60)*

#### 13. Common pncA Mutation Codon

Codon **94** (e.g., Asp94Gly/Asn) is frequently mutated in PZA resistance. *(Section 3*.*5, Page 36)*

#### 14. tlyA and Capreomycin Resistance

LoF mutations in *tlyA* (ribosomal RNA methyltransferase) disrupt capreomycin activation, conferring resistance. *(Section 3*.*10, Page 64)*

#### 15. Compensatory Mutation in rpoC

**rpoC Pro432Leu** is a compensatory mutation linked to rifampicin-resistant strains. *(Table 6, Page 34)*

#### 16. Bedaquiline Resistance Genes

*Rv0678* and *pepQ* are key genes. LoF in these genes is associated with resistance. *(Section 3*.*7, Page 52; Table 1, Page 16)*

#### 17. katG Trp328Leu Significance

Group 1 mutation conferring high-level INH resistance. *(Table 7, Page 37; Section 3*.*3, Page 26)*

#### 18. gyrA Codon 94 Impact

Mutations (e.g., Asp94Gly) confer high-level fluoroquinolone resistance. *(Section 3*.*6, Page 48; Table 1, Page 16)*

#### 19. pncA Leu116Pro and PZA Resistance

Yes. Leu116Pro is a Group 1 mutation associated with PZA resistance. *(Section 3*.*5, Page 36)*

#### 20. embB M306V in EMB Resistance

**embB Met306Val** is a major resistance mutation (Group 1), altering ethambutol target binding. *(Section 3*.*4, Page 33)*

#### 21. rpoB S450L Molecular Mechanism

Ser450Leu in the RRDR disrupts rifampicin binding to RNA polymerase. *(Section 3*.*2, Page 29; Table 6, Page 30)*

#### 22. rpsL Streptomycin Resistance Mutation

**rpsL Lys43Arg** is a common Group 1 mutation. *(Section 3*.*11, Page 69)*

#### 23. embA Regulatory Region in EMB Resistance

Yes. Promoter mutations (e.g., −16A>G) upregulate *embCAB* operon, causing resistance. *(Section 3*.*4, Page 33)*

#### 24. Non-RRDR rpoB Mutations

**rpoB Val170Phe** and **Ile491Phe** outside RRDR are Group 1 resistance mutations. *(Table 6, Page 30; Section 3*.*2, Page 29)*

#### 25. pepQ Mutation for Delamanid Resistance

**pepQ Gly20**\* (LoF) is a Group 2 mutation. *(Section 3*.*9, Page 62)*

#### 26. mmpL5 Interaction with Rv0678

**mmpL5 frameshift mutations** (e.g., Trp20*) negate Rv0678-mediated BDQ resistance via epistasis. *(Section 3*.*7, Page 52)*

#### 27. pncA Duplications

No specific duplications mentioned, but frameshifts/indels (e.g., Phe106fs) are documented. *(Section 3*.*5, Page 36)*

#### 28. katG S315T vs. inhA −15C>T

- **katG S315T**: High-level INH resistance (Group 1).
- **inhA −15C>T**: Low-level resistance (Group 1), cross-resistant to ethionamide. *(Section 3*.*3, Page 26; Table 7, Page 37)*

#### 29. Frameshift vs. Missense in pncA

Both are Group 1/2 for PZA resistance. Frameshifts (LoF) abolish PncA activity, while missense may reduce enzyme function. *(Section 3*.*5, Page 36)*

#### 30. Large katG Deletions

Yes. LoF deletions (e.g., katG Met1?) are Group 1 for INH resistance. *(Table 7, Page 37)*

**References:** All answers derived from the provided document pages and sections as cited. Specific tables and sections are noted for verification.

## DEEPSEEK - PROMPT 3

1. **katG_p.Lys154fs (INH):**
  ∘ **Group 2 (Assoc w R-interim)**
  ∘ **Drug:** Isoniazid
  ∘ **Rule:** LoF mutations in *katG* are classified as Group 2 per the INH-specific additional grading rule (Table 1). Frameshift mutations are considered LoF.
2. **pncA_p.Arg140Ala (PZA):**
  ∘ **Group 3 (Uncertain significance)**
  ∘ **Drug:** Pyrazinamide
  ∘ **Rule:** Missense mutations in *pncA* not explicitly graded in the catalogue default to Group 3. Only LoF mutations (nonsense/frameshift) trigger the Group 2 rule (Section 2.1, Table 1).
3. **Rv0678_p.Met1? (BDQ/CFZ):**
  ∘ **Group 2 (Assoc w R-interim)**
  ∘ **Drugs:** Bedaquiline and clofazimine
  ∘ **Rule:** Start-loss mutations are LoF. *Rv0678* LoF mutations confer cross-resistance to BDQ/CFZ (Section 3.7, Table 1).
4. **fbiC deletion (DLM/PMD):**
  ∘ **Group 2 (Assoc w R-interim)**
  ∘ **Drugs:** Delamanid and pretomanid
  ∘ **Rule:** Complete gene deletions in *fbiC* are LoF, classified as Group 2 per DLM/PMD-specific rules (Section 3.9, Table 1).
5. **rpoB_p.Gln432Arg (RIF):**
  ∘ **Group 2 (Assoc w R-interim)**
  ∘ **Drug:** Rifampicin
  ∘ **Rule:** Non-synonymous mutations in the RRDR (*rpoB* codons 426–452) are automatically Group 2 (Section 3.2, Table 1).
6. **fabG1_c.-8T>A (INH/ETO):**
  ∘ **Group 2 (Assoc w R-interim)**
  ∘ **Drugs:** Isoniazid and ethionamide
  ∘ **Rule:** *fabG1-inhA* promoter mutations confer cross-resistance to INH and ETO (Section 3.3, Table 1).
7. **gyrA_p.Gly88Ala (FQs):**
  ∘ **Group 2 (Assoc w R-interim)**
  ∘ **Drugs:** Levofloxacin and moxifloxacin
  ∘ **Rule:** Non-synonymous *gyrA* mutations are Group 2. Gly88Ala is not a high-level MFX mutation (unlike Gly88Cys), so it confers low-level resistance (Section 3.6, Table 1).
8. **eis_c.-14C>T + eis_p.Trp50fs (AMK):**
  ∘ **Group 5 (Not assoc w R)**
  ∘ **Drug:** Amikacin
  ∘ **Rule:** *eis* promoter mutations (−14C>T) are negated by coding LoF mutations (Trp50fs) due to epistasis (Section 3.10, Table 1).
9. **gyrB_p.Gly520Gly (FQs):**
  ∘ **Group 4 (Not assoc w R-interim)**
  ∘ **Drugs:** Fluoroquinolones
  ∘ **Rule:** Synonymous mutations are classified as Group 4 per the general additional grading rule (Section 2.1, Fig. 1).
10. **Rv0678_p.Gly121Arg + mmpL5_LoF (BDQ):**
  ∘ **Group 5 (Not assoc w R)**
  ∘ **Drug:** Bedaquiline
  ∘ **Rule:** *Rv0678* mutations (Gly121Arg) are epistatically suppressed by *mmpL5* LoF, preventing resistance (Section 3.7, Table 1).

## CHATGPT - PROMPT 1

Based solely on the contents of the *WHO 2023 Catalogue of Mutations in Mycobacterium tuberculosis*, here are the answers to your questions:

1. **Describe the different groups used to classify mutations in the catalogue**. Mutations are classified into five groups based on the statistical strength of their association with drug resistance:
  - **Group 1:** Associated with resistance (Assoc w R)
  - **Group 2:** Associated with resistance – interim (Assoc w R–interim)
  - **Group 3:** Uncertain significance
  - **Group 4:** Not associated with resistance – interim (Not assoc w R–interim)
  - **Group 5:** Not associated with resistance (Not assoc w R)
  Groups 1 and 2 are markers of phenotypic resistance; Groups 4 and 5 are not. Group 3 mutations are of uncertain significance due to insufficient or conflicting evidence.
2. **How are mutations graded in the catalogue?** Mutations are graded using criteria involving:
  - Allele frequency (*≥*75% in at least one isolate)
  - Valid phenotypic DST (pDST) result
  - SOLO algorithm statistical metrics (e.g., odds ratio, PPV|SOLO thresholds)
  The grading uses a five-tier system (Groups 1–5) based on these criteria and additional grading rules (both general and drug-specific).
3. **Explain what is meant by “Uncertain Significance” mutations**. Group 3 mutations (“Uncertain significance”) do not meet the criteria for inclusion in Groups 1, 2, 4, or 5. Their effect on resistance is unclear due to insufficient statistical association or lack of phenotypic data.
4. **How do you interpret a mutation listed as “Assoc w R-interim”?** “Assoc w R–interim” (Group 2) mutations show a statistically significant but less robust association with resistance than Group 1 mutations. They are still interpreted as **clinically relevant markers of phenotypic resistance**, especially when supported by additional grading rules.
5. **Explain the concept of “loss-of-function” (LoF) mutations and their relevance to drug resistance**. LoF mutations result in the disruption of protein function, typically via frameshifts, premature stop codons, or deletions. In the catalogue, LoF mutations in specific genes (e.g., *katG, pncA, Rv0678*) are associated with resistance and are graded under drug-specific rules as Group 2 unless otherwise stated.
6. **Does the catalogue contain mutations that are not associated with drug resistance?** Yes. Group 4 and 5 mutations are **not associated** with resistance. Group 4 includes variants with intermediate statistical evidence against resistance, and Group 5 includes those deemed neutral, often silent or previously masked by the algorithm.
7. **What is the most common resistance-conferring mutation for rifampicin?** The **rpoB Ser450Leu** (S450L) mutation is the most frequent and well-established rifampicin resistance mutation.
8. **Identify the nucleotide change associated with the most frequent inhA promoter mutation linked to isoniazid resistance**. The **fabG1 −15C>T** mutation (also known as inhA −777C>T) is the most frequent promoter mutation linked to isoniazid resistance.
9. **What is the role of “epistasis” in interpreting resistance to bedaquiline and clofazimine?** Epistasis refers to **interactions between mutations** that affect the resistance phenotype. For bedaquiline and clofazimine:
  - *Rv0678* mutations confer resistance
  - If *Rv0678* mutations co-occur with **LoF mutations in mmpL5**, resistance is negated (epistasis cancels the effect).
10. **Identify a mutation in the Rv0678 gene that is associated with bedaquiline resistance**. One example is **Rv0678_p.Ile67Ser**, which is listed as Group 2 (Assoc w R–interim) for bedaquiline resistance.
11. **Are insertions or deletions (indels) in the pncA gene known to cause pyrazinamide resistance?** Yes, LoF mutations in *pncA*, including indels, are associated with pyrazinamide resistance. These are generally graded under drug-specific rules and considered Group 2 unless otherwise excluded.
12. **Locate a mutation that is associated with resistance to linezolid in the rrl gene**. Information not found in document snippet. Please refer to the full mutation table for *rrl* mutations under linezolid.
13. **What is the codon position of the most common pyrazinamide resistance mutation in the pncA gene?** Information not found in document snippet. The full mutation table or supplementary Excel file is required.
14. **How does the tlyA gene contribute to capreomycin resistance?** LoF mutations in *tlyA* are associated with capreomycin resistance. These are classified under additional grading rules as Group 2 mutations.
15. **Provide an example of a compensatory mutation in the rpoC gene that is found in rifampicin-resistant strains**. Information not found in document snippet. Please refer to the *rpoC* section in the catalogue.
16. **What genes are involved in bedaquiline resistance?**
  - *Rv0678*
  - *atpE*
  - *pepQ*
  - Epistatic interactions with *mmpL5* also influence resistance.
17. **A Trp328Leu mutation in katG has been found in a sample. What does it mean?** Information not found in document snippet. Please check the *katG* entry in the full mutation table.
18. **Explain the potential impact of a mutation at codon 94 of gyrA**. Mutations at *gyrA codon 94* (e.g., Asp94Gly, Asp94Asn) are associated with **high-level resistance to moxifloxacin** and likely also to levofloxacin.
19. **Does the Leu116Pro mutation in pncA confer resistance to pyrazinamide?** Information not found in document snippet. Requires lookup in the mutation table.
20. **What is the significance of the embB M306V mutation in ethambutol resistance?** Information not found in document snippet. Check the *embB* section in the full catalogue.
21. **How does the rpoB S450L mutation lead to rifampicin resistance at the molecular level?** Information not found in document snippet. The catalogue states it is associated with resistance but does not detail the **molecular mechanism**.
22. **Find an example of a mutation in the rpsL gene that is associated with streptomycin resistance**. Information not found in document snippet. Please refer to the *rpsL* entries in the mutation tables.
23. **Is the embA regulatory region involved in ethambutol resistance?** Yes. Mutations in the *embA* upstream region (like *fabG1* for *inhA*) are included in resistance analysis based on their regulatory impact.
24. **Are there any mutations outside the rifampicin resistance-determining region (RRDR) of rpoB that contribute to resistance?** Yes, although the RRDR is the primary focus, other mutations (e.g., at codon 533) outside RRDR have been reported and interpreted with caution.
25. **Find an example of a mutation in the pepQ gene that has been linked to resistance against delamanid**. *pepQ* LoF mutations are associated with resistance to **both bedaquiline and clofazimine**, but delamanid is not explicitly linked in the retrieved snippet.
26. **Identify a mutation in the mmpL5 gene that interacts with Rv0678 mutations in the context of bedaquiline resistance**. LoF mutations in *mmpL5* negate resistance conferred by *Rv0678* mutations (epistasis).
27. **Does the pncA gene contain any known duplications that contribute to pyrazinamide resistance?** Information not found in document snippet. Please refer to the *pncA* mutation list.
28. **Compare the impact of the katG S315T versus an inhA −15C>T on isoniazid resistance**.
  - **katG S315T** *→* High-level INH resistance (Group 1)
  - **inhA −15C>T (fabG1)** *→* Low-level INH resistance (Group 2)
  They may act synergistically if co-occurring.
29. **How does a frameshift mutation in pncA compare to a missense SNP in pncA in terms of pyrazinamide resistance?** Frameshift mutations (LoF) are more likely to confer resistance and are graded via the LoF rule (Group 2). Some missense SNPs may also confer resistance but are graded based on statistical evidence.
30. **Are there any documented instances of large deletions in the katG gene, and do they impact isoniazid resistance?** Yes. Large deletions and other LoF mutations in *katG* are associated with **high-level isoniazid resistance** and are considered Group 2 under additional grading rules.

## CHATGPT - PROMPT 3

Based on the WHO 2023 Mutation Catalogue and the interpretation framework provided in Section 2.1 and Table 1, here are detailed responses to your questions:

1. **katG_p.Lys154fs — Interpretation for Isoniazid Resistance**
  - **Type:** Novel frameshift mutation in katG.
  - **Rule Applied:** *Additional grading rule* for INH: **Loss-of-function (LoF) in katG *→* Group 2 (Assoc w R–interim)**.
  - **Interpretation:** Predicted **high-level resistance to isoniazid**.
  - **Confidence Group: Group 2**
  - **Justification:** Table 1 (p. 6) — LoF mutations in katG are assumed to confer high-level INH resistance and are classified under Group 2.
2. **pncA_p.Arg140Ala — Interpretation for Pyrazinamide Susceptibility**
  - **Type:** Nonsense mutation in pncA.
  - **Rule Applied:** If this is a true nonsense mutation (although the name suggests missense), and is LoF:
    ∘ **LoF in pncA *→* Group 2 (Assoc w R–interim)** for pyrazinamide.
  - **Interpretation: Resistant to pyrazinamide**
  - **Confidence Group: Group 2**, assuming it is LoF.
  - **Justification:** Table 1 — LoF in pncA assumed to confer PZA resistance.
3. **Rv0678_p.Met1? — Interpretation for Drug Resistance**
  - **Type:** Start-loss mutation in Rv0678.
  - **Rule Applied:** LoF in Rv0678 *→* Group 2 (Assoc w R–interim) for **bedaquiline and clofazimine**.
  - **Interpretation: Associated with resistance to bedaquiline and clofazimine**.
  - **Confidence Group: Group 2**
  - **Justification:** Table 1 — LoF in Rv0678 is considered cross-resistance to BDQ and CFZ.
4. **fbiC Deletion — Effect on Delamanid Resistance**
  - **Type:** LoF via deletion of entire gene.
  - **Rule Applied:** LoF in fbiC *→* Group 2 (Assoc w R–interim) for **delamanid**.
  - **Interpretation: Predicted resistance to delamanid (and pretomanid)**.
  - **Confidence Group: Group 2**
  - **Justification:** Table 1 — LoF in fbiC assumed to confer resistance to DLM and PMD.
5. **rpoB_p.Gln432Arg — Classification for Rifampicin Resistance**
  - **Type:** Missense mutation in the RRDR of rpoB.
  - **Found in Catalogue:** Yes.
  - **Group: Group 1 (Assoc w R)**
  - **Interpretation: Rifampicin resistant**.
  - **Confidence Group: Group 1**
  - **Justification:** Found in abridged mutation tables; meets Group 1 criteria (p. 16, Fig. 4).
6. **fabG1_c.-8T>A — Implications for INH and ETH Resistance**
  - **Type:** Promoter mutation.
  - **Rule Applied:** fabG1-inhA mutations confer low-level INH resistance and cross-resistance to ethionamide.
  - **Interpretation:**
    ∘ **Low-level isoniazid resistance**
    ∘ **Cross-resistance to ethionamide**
  - **Confidence Group: Group 2 (Assoc w R–interim)**
  - **Justification:** Table 1 — Listed among mutations with assumed cross-resistance.
7. **gyrA_p.Gly88Ala — Interpretation for Fluoroquinolones**
  - **Type:** Non-synonymous mutation in gyrA.
  - **Found in Catalogue:** Yes.
  - **Confidence Group: Group 1 (Assoc w R)**
  - **Interpretation:**
    ∘ **Levofloxacin and moxifloxacin resistance**
    ∘ **High-level MFX resistance** (per Table 1)
  - **Justification:** Table 1 — gyrA Gly88 mutations are assumed to confer high-level resistance to moxifloxacin.
8. **eis_c.-14C>T + eis_p.Trp50fs — Interpretation for Amikacin Resistance**
  - **Type:** Combination of promoter and coding LoF mutation.
  - **Rule Applied:** Epistasis: eis promoter mutation cannot confer resistance if genetically linked with a coding LoF variant.
  - **Interpretation: Not resistant to amikacin**.
  - **Confidence Group: Not a resistance marker (Group 4/5)**.
  - **Justification:** Table 1 — Epistasis rule for AMK/KAN.
9. **gyrB_p.Gly520Gly — Synonymous Mutation**
  - **Type:** Novel synonymous mutation.
  - **Rule Applied:** General rule — all novel silent mutations are Group 4.
  - **Interpretation: Not associated with resistance**
  - **Confidence Group: Group 4 (Not assoc w R–interim)**
  - **Justification:** Section 2.1 and Fig. 1 workflow — novel silent variants are Group 4.
10. **Rv0678_p.Gly121Arg + mmpL5_LoF — Interpretation for Bedaquiline Resistance**
  - **Type:** Rv0678 mutation plus mmpL5 LoF.
  - **Rule Applied:** Epistasis: Rv0678 mutations **do not confer resistance** if linked to LoF in mmpL5.
  - **Interpretation: Not resistant to bedaquiline**
  - **Confidence Group: Not a resistance marker**
  - **Justification:** Table 1 — Rv0678 mutations require functional mmpL5 for resistance.

## PERPLEXITY - PROMPT 1

1. The catalogue classifies mutations into four main groups:
  - Associated with resistance (Assoc w R)
  - Associated with resistance–interim (Assoc w R–interim)
  - Not associated with resistance (Not assoc w R)
  - Not associated with resistance–interim (Not assoc w R–interim)5
2. Mutations are graded in the catalogue using a confidence grading system based on criteria such as:
  - Frequency of occurrence in resistant isolates
  - Absence in susceptible isolates
  - Experimental or clinical evidence
  - Biological plausibility The grades range from “Uncertain significance” to “High confidence”[97-101]
3. “Uncertain significance” mutations are those that do not meet the criteria for other confidence grades. They may have conflicting evidence or insufficient data to determine their association with resistance100
4. “Assoc w R-interim” indicates a mutation is provisionally associated with resistance, but requires additional evidence to confirm this association5
5. Loss-of-function (LoF) mutations are those that disrupt gene function, such as nonsense mutations, frameshift indels, or large deletions. For some drugs like pyrazinamide, LoF mutations in the pncA gene are a major mechanism of resistance36
6. Yes, the catalogue contains mutations that are not associated with drug resistance. These are classified as “Not assoc w R” or “Not assoc w R–interim”5
7. The most common resistance-conferring mutation for rifampicin is rpoB S450L19
8. The most frequent inhA promoter mutation linked to isoniazid resistance is c-15t26
9. Epistasis refers to interactions between mutations that can affect drug resistance. For bedaquiline and clofazimine, mutations in Rv0678 can interact with mutations in mmpL5 to confer resistance52
10. The L117P mutation in the Rv0678 gene is associated with bedaquiline resistance52
11. Yes, insertions and deletions (indels) in the pncA gene are known to cause pyrazinamide resistance36
12. The G2270T/C mutation in the rrl gene is associated with resistance to linezolid60
13. The most common pyrazinamide resistance mutation in the pncA gene is at codon 15936
14. Information about tlyA gene’s contribution to capreomycin resistance is not found in the provided document.
15. The I491F mutation in rpoC is an example of a compensatory mutation found in rifampicin-resistant strains19
16. The genes involved in bedaquiline resistance are Rv0678, atpE, and pepQ52
17. A Trp328Leu mutation in katG is associated with isoniazid resistance26
18. Mutations at codon 94 of gyrA are associated with resistance to fluoroquinolones like levofloxacin and moxifloxacin48
19. Yes, the Leu116Pro mutation in pncA confers resistance to pyrazinamide36
20. The embB M306V mutation is associated with ethambutol resistance33
21. The rpoB S450L mutation leads to rifampicin resistance by altering the structure of the RNA polymerase beta subunit, which reduces rifampicin binding19
22. The K43R mutation in the rpsL gene is associated with streptomycin resistance
23. Yes, the embA regulatory region is involved in ethambutol resistance33
24. Yes, there are mutations outside the RRDR of rpoB that contribute to resistance, such as I491F19
25. The D26fs mutation in the pepQ gene has been linked to resistance against delamanid
26. The G448S mutation in the mmpL5 gene interacts with Rv0678 mutations in the context of bedaquiline resistance52
27. Yes, the pncA gene contains known duplications that contribute to pyrazinamide resistance36
28. The katG S315T mutation generally confers high-level isoniazid resistance, while the inhA −15C>T mutation typically confers low-level resistance26
29. Frameshift mutations in pncA generally lead to complete loss of function and high-level pyrazinamide resistance, while missense SNPs may have variable effects depending on their location and impact on protein function36
30. Yes, there are documented instances of large deletions in the katG gene, which can impact isoniazid resistance by causing loss of function of the KatG enzyme

## PERPLEXITY - PROMPT 3

Here is the analysis of each mutation based on the WHO Catalogue of Mutations (2nd edition):

1. katG_p.Lys154fs (Isoniazid)
  - Confidence Group: 1
  - Drug: Isoniazid
  - Rule: Frameshift mutations in katG are classified as “Associated with resistance” (Assoc w R) due to LoF (loss-of-function) effects (Section 3.3). katG mutations account for ~60-90% of isoniazid resistance1.
2. pncA_p.Arg140Ala (Pyrazinamide)
  - Confidence Group: 1
  - Drug: Pyrazinamide
  - Rule: All non-synonymous pncA mutations (including missense/nonsense) outside the promoter are Assoc w R (Section 3.5). This excludes synonymous variants and promoter mutations1.
3. Rv0678_p.Met1? (Bedaquiline/Clofazimine)
  - Confidence Group: 1
  - Drugs: Bedaquiline, Clofazimine
  - Rule: Start-loss mutations in Rv0678 (mmpR) are Assoc w R due to disrupted repression of mmpS5-mmpL5 efflux pump (Section 3.7). Requires reporting for both drugs1.
4. fbiC full deletion (Delamanid)
  - Confidence Group: 1
  - Drug: Delamanid
  - Rule: Complete deletion of fbiC (required for F420 cofactor biosynthesis) is Assoc w R. FbiC LoF prevents prodrug activation (Section 3.9)1.
5. rpoB_p.Gln432Arg (Rifampicin)
  - Confidence Group: 1
  - Drug: Rifampicin
  - Rule: Non-synonymous RRDR (rifampicin resistance-determining region) mutations are Assoc w R (Section 3.2). Gln432Arg is a known resistance mutation1.
6. fabG1_c.-8T>A (Isoniazid/Ethionamide)
  - Confidence Group: 1
  - Drugs: Isoniazid, Ethionamide
  - Rule: fabG1 promoter mutations at positions −15 to −8 are Assoc w R for both drugs (Section 3.3, 3.12). Affects inhA expression1.
7. gyrA_p.Gly88Ala (Fluoroquinolones)
  - Confidence Group: 1
  - Drugs: Levofloxacin, Moxifloxacin
  - Rule: gyrA mutations at codons 88-94 are Assoc w R (Section 3.6). Gly88Ala is a recognized resistance mutation1.
8. eis_c.-14C>T + eis_p.Trp50fs (Amikacin)
  - Confidence Group: 5
  - Drug: Amikacin
  - Rule: The frameshift (Trp50fs) causes LoF, overriding the resistance-associated promoter mutation (−14C>T). Catalogued as “Not assoc w R” when disruptive mutations occur (Section 3.10)1.
9. gyrB_p.Gly520Gly (Fluoroquinolones)
  - Confidence Group: 5
  - Drugs: Fluoroquinolones
  - Rule: Synonymous mutations outside known resistance-associated regions are “Not assoc w R” (Section 3.6). No amino acid change1.
10. Rv0678_p.Gly121Arg + mmpL5_LoF (Bedaquiline)
  - Confidence Group: 5
  - Drug: Bedaquiline
  - Rule: mmpL5 LoF mutations negate resistance from Rv0678 variants (Section 3.7). Functional mmpL5 is required for efflux-mediated resistance1.

Citations refer to specific mutation tables and interpretation rules in Sections 3.2-3.12 of the catalogue1.

